# Generating Biomedical Knowledge Graphs from Knowledge Bases, Registries, and Multiomic Data

**DOI:** 10.1101/2024.11.14.623648

**Authors:** Guangrong Qin, Kamileh Narsinh, Qi Wei, Jared C. Roach, Arpita Joshi, Skye L. Goetz, Sierra T. Moxon, Matthew H. Brush, Colleen Xu, Yao Yao, Amy K. Glen, Evan D. Morris, Alexandra Ralevski, Ryan Roper, Basazin Belhu, Yue Zhang, Ilya Shmulevich, Jennifer Hadlock, Gwênlyn Glusman

**Author notes:** Correspondence, Gwênlyn Glusman, Institute for Systems Biology, 401 Terry Ave N, Seattle, WA 98109, USA. co-first authors (Guangrong Qin, Kamileh Narsinh). deceased (Ilya Shmulevich).

## Abstract

As large clinical and multiomics datasets and knowledge resources accumulate, they need to be transformed into computable and actionable information to support automated reasoning. These datasets range from laboratory experiment results to electronic health records (EHRs). Barriers to accessibility and sharing of such datasets include diversity of content, size and privacy. Effective transformation of data into information requires harmonization of stakeholder goals, implementation, enforcement of standards regarding quality and completeness, and availability of resources for maintenance and updates. Systems such as the Biomedical Data Translator leverage knowledge graphs (KGs), structured and machine learning readable knowledge representation, to encode knowledge extracted through inference. We focus here on the transformation of data from multiomics datasets and EHRs into compact knowledge, represented in a KG data structure. We demonstrate this data transformation in the context of the Translator ecosystem, including clinical trials, drug approvals, cancer, wellness, and EHR data. These transformations preserve individual privacy. We provide access to the five resulting KGs through the Translator framework. We show examples of biomedical research questions supported by our KGs, and discuss issues arising from extracting biomedical knowledge from multiomics data.

## 1. Introduction

Knowledge Graphs (KGs) are abstract representations of available knowledge, encoding entities as ‘nodes’, and the established relationships between these entities as ‘edges’, supported by attributes denoting evidence, provenance, and confidence. KGs are networks that can be described atomically by triples of ‘subject’, ‘predicate’, and ‘object’—also known as ‘head entity’, ‘relationship’, ‘tail entity’ or ‘source node’, ‘relation’, ‘target node’. KGs are increasingly used to model biomedical research data and to inform better predictions^1–5^. These models capture prior data and can be used in conjunction with reasoning algorithms to answer questions about these data. More formally, hypotheses can be tested by querying a KG. The Biomedical Data Translator program (‘Translator’), funded by the National Center for Advancing Translational Sciences (NCATS), aims to facilitate the transformation of data resulting from basic science projects into clinically actionable knowledge to drive research innovations^6–10^. Translator is being developed by a consortium of teams from two dozen institutions^6^, that are constructing both KGs and developing autonomous relay agents (ARAs) to implement integrative reasoning algorithms on KGs. The Translator is a federated system where KGs are encoded in the same standard^10^ and exposed through an application programming interface (API) following the Translator standard (TRAPI)^11^.

The fundamental basis to extract KGs that can be used in the Translator system is the massive data and knowledge generated in the biomedical community. Data are units of information, often numeric, gathered through observation or measurement that represent real-world phenomena^12^. In the context of the KGs discussed here, knowledge is a human belief or machine assertion justified by evidence that is likely to be true^13^. Knowledge can be generated by combining data and reasoning^14^. The sources of knowledge introduced into Translator can be classified into three main categories: **(1) preexisting knowledge easily and unambiguously extracted from reliable sources, (2) preexisting knowledge extracted—ambiguously or uncertainly—from sources of varying reliability, and (3) new knowledge derived from computational analysis of large datasets**. Reliable sources of the first kind are typically well curated knowledge bases, but may include other publications and repositories. Sources of the second kind include knowledge text-mined from scientific publications, and from registries with limited curation of data provided by researchers. Excitingly, new knowledge lies latent (yet to be perceived by humans) in large collections of clinical and scientific data; we discuss here several approaches to unearth this knowledge and store the results in accessible KGs, with particular focus on analysis of multiomic datasets and electronic health records (EHRs).

KGs extracted from different data resources, including multiomics data from cancer and wellness, large scale real world clinical data, and clinical trial data, provide rich resources to understand and explain the relationship among diseases, genes and drugs in different contexts. Despite the different resources each KG consumed, we present a general pipeline—in the context of Translator—for production of KGs from multiomic data and knowledge resources. We describe five KGs that we make accessible through the Translator infrastructure. <u>BigGIM-DrugResponse KG</u> is derived from large multiomics data, *ex vivo* drug response data, and accumulated knowledge resources to capture relationships among diseases, molecular features, and drugs or chemicals. <u>Clinical Trials KG</u> and <u>Drug Approvals KG</u> encode relationships between interventions and conditions, as mined from three registries: clinicaltrials.gov, DailyMed^15^, and the FDA’s Adverse Event Reporting System (FAERS)^16^. <u>Clinical Connections KG</u> is derived from massive electronic medical record databases that connect concepts (such as diseases) with other clinical concepts such as outcomes. Finally, <u>Wellness Multiomics KG</u> is derived from a large longitudinal healthy cohort and encodes relationships between omics analytes and clinical variables.

### Reasoning from Data to Knowledge

We use several automated epistemological approaches from data science to automatically reason upon large multiomic and EHR datasets, and so produce knowledge from these data. Data science^17^ encompasses the creation of principles and algorithms that enable extraction of information and knowledge from data^18,19^. Humans use multiple categories of reasoning to extract knowledge from data^20^. In bioscience, these categories have been termed “Hill’s criteria”^21^, and include statistical significance, effect size, analogy, specificity, time dependence, dose-response, consistency, reversibility, validation, coherence, and biological plausibility. A data-science perspective is to use these categories systematically in a framework to extract useful knowledge from data. For purposes of machine-automatable KG construction designed for wide use in the biological community, we generally require that our data-science framework be understandable, explainable, and reproducible—with solid provenance. Therefore we focus on statistics and explainable machine learning for edges based on newly generated knowledge in the KGs reported here. We avoid inference based on categories of reasoning that are currently better implemented by humans such as analogy and biological plausibility. However, one of the main purposes of these KGs is to facilitate exactly those kinds of reasoning, making it easier for humans to go beyond basic retrieval of facts. A utility of these and other KGs is to provide a compilation of pre-existing knowledge that humans can use for reasoning based on Hill’s criteria, Bayesian inference, and/or other preferred epistemological approaches. KGs are a structured form of prior knowledge.

Automated data mining and statistical analysis largely create knowledge by first identifying correlations or associations among concepts. Data mining may reveal unexpected relationships—fostering new conceptualizations and fostering innovative knowledge^22^.

KGs also provide more mundane benefits that are profoundly important for biomedical research. KGs provide an efficient, computer-readable structure for sharing data. In some cases, this method of sharing enables private (e.g., HIPAA) data to be shared as public knowledge by aggregating, reasoning upon, or de-identifying raw data. KGs provide pre-computed analyses (e.g., correlations) that are likely to be interesting and/or would have to be performed by many users in parallel downstream of the data; it makes sense to pre-compute these analyses for efficiency and accessibility. KGs also can be comprehensive, losing little to no information from a data system and therefore avoid self-imposed limitations that might arise from use of a more lossy or limited (often manually curated) data structure.

## 2. Results

### 2.1. Overview

We analyzed and curated evidence from EHR data, multiomics data from normal tissue^23^ & cancers^24,25^, a wellness cohort^26^, and public knowledge resources, and generated KGs representing connections observed in the data (**Table 1**). These KGs result from a workflow—described in Methods—that transforms multiomics data to knowledge including (1) collection of data resources, (2) data pre-processing; (3) statistical modeling, (4) standardization using the Biolink Model^10^; and (5) implementation of APIs to expose the KGs to users and the Translator ecosystem (**Figure 1**). We constructed five KGs: BigGIM-Drug Response KG, Wellness KG, Clinical Trials KG, Drug Approvals KG, and Clinical Connections KG.

**Table 1.** Characteristics of the five Knowledge Graphs.

| Knowledge Graph | Relationship Type | KG Source Data | Node Categories (and CURIE prefixes) | Biolink Predicates |
| --- | --- | --- | --- | --- |
| BigGIM - Drug Response | Gene expression to gene expression | GTEX; TCGA | Gene expression (NCBIGene) | positively/negatively correlated with |
|  | Gene to gene | BioGrid | Gene (NCBIGene) | genetically interacts with |
|  | Protein to protein | BioGrid, HuRI | Gene (NCBIGene) | physically interacts with |
|  | Gene mutation to drug response | GDSC | Gene mutation (NCBIGene), Drug/Chemical (PUBCHEM) | associated with sensitivity/resistance to |
|  | Gene expression to drug response | GDSC | Gene expression (NCBIGene), Drug/Chemical (PUBCHEM) | associated with sensitivity/resistance to |
|  | Drug to target | Publicly available knowledge resources (Drug Central, TTD, Pubmed) | Gene (NCBIGene), Drug (PUBCHEM) | physically interacts with |
|  | Cell to gene | CellMarker | Gene (NCBIGene) Cell (CL) | expressed in |
|  | TCGA Mutation Frequency | TCGA | Disease (MONDO), Gene (NCBIGene) | gene associated with condition, has biomarker |
| Wellness Multiomics | Statistical association | ISB Wellness data on clinical labs, proteins, metabolites | ClinicalFinding (LOINC, NCIT, MESH), Protein (UniProtKB), SmallMolecule (PUBCHEM, HMDB, EFO, CAS, KEGG, CHEBI, UMLS) | correlated with |
| Clinical Connections | Risk factors for chronic diseases | Providence Health & Services EHRs | ChemicalEntity (CHEBI, RXCUI, UNII), Disease (MONDO, SNOMEDCT), PhenotypicFeature (HP, NCIT) | associated with increased/decreased likelihood of |
| Clinical Trials | Intervention to condition | ClinicalTrials.gov | Disease (UMLS, MONDO, EFO, DOID, NCIT, OMIM, MESH), PhenotypicFeature (HP, UMLS, NCIT, EFO, MESH), SmallMolecule/MolecularMixture (CHEBI, PUBCHEM, ChEMBL), | in clinical trials for, treats |
|  |  |  | ChemicalEntity (UNII) |  |
| Drug Approvals | Intervention to condition | DailyMed, FAERS | Disease (UMLS, MONDO, EFO, DOID, NCIT, OMIM, MESH),<br>PhenotypicFeature (HP, UMLS, NCIT, EFO, MESH),<br>SmallMolecule/MolecularMixture (CHEBI, PUBCHEM, ChEMBL),<br>ChemicalEntity (UNII) | treats,<br>applied to treat |

**Figure 1.**
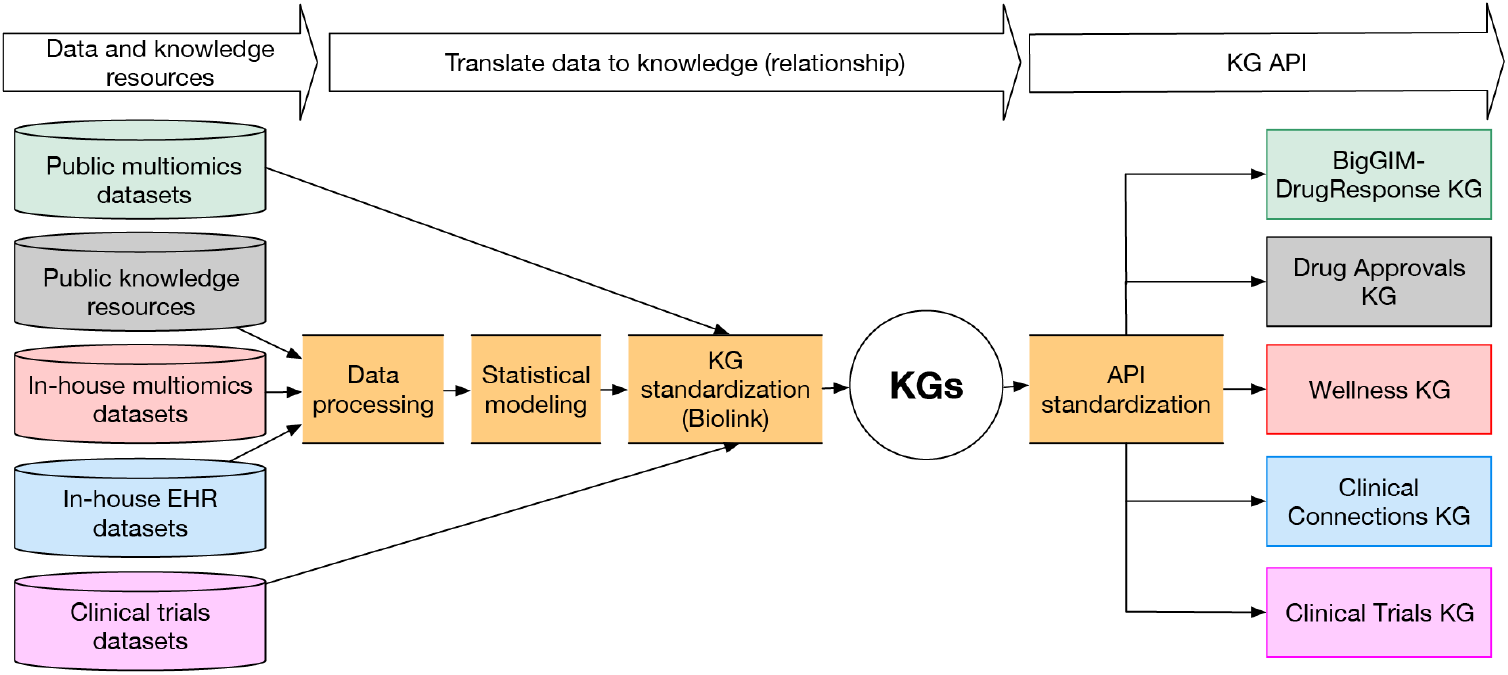
Workflow Overview. Conceptual overview of the pipeline for translating multiomics & EHR data and knowledge resources to Translator standard KGs and Application Programming Interfaces (APIs). KGs are standardized using the Biolink Model to facilitate reasoning leveraging multiple KGs.

The metagraph for the merged KGs (**Figure 2**, and detailed in <u>Supplementary Table 1</u>) indicates the general categories of concepts and relationships in each KG, and not only summarizes the value of each individual KG, but also provides a strategic map of how these concepts interconnect. This map can guide application of a federated system of KGs, such as Translator, for specific use cases.

**Figure 2.**
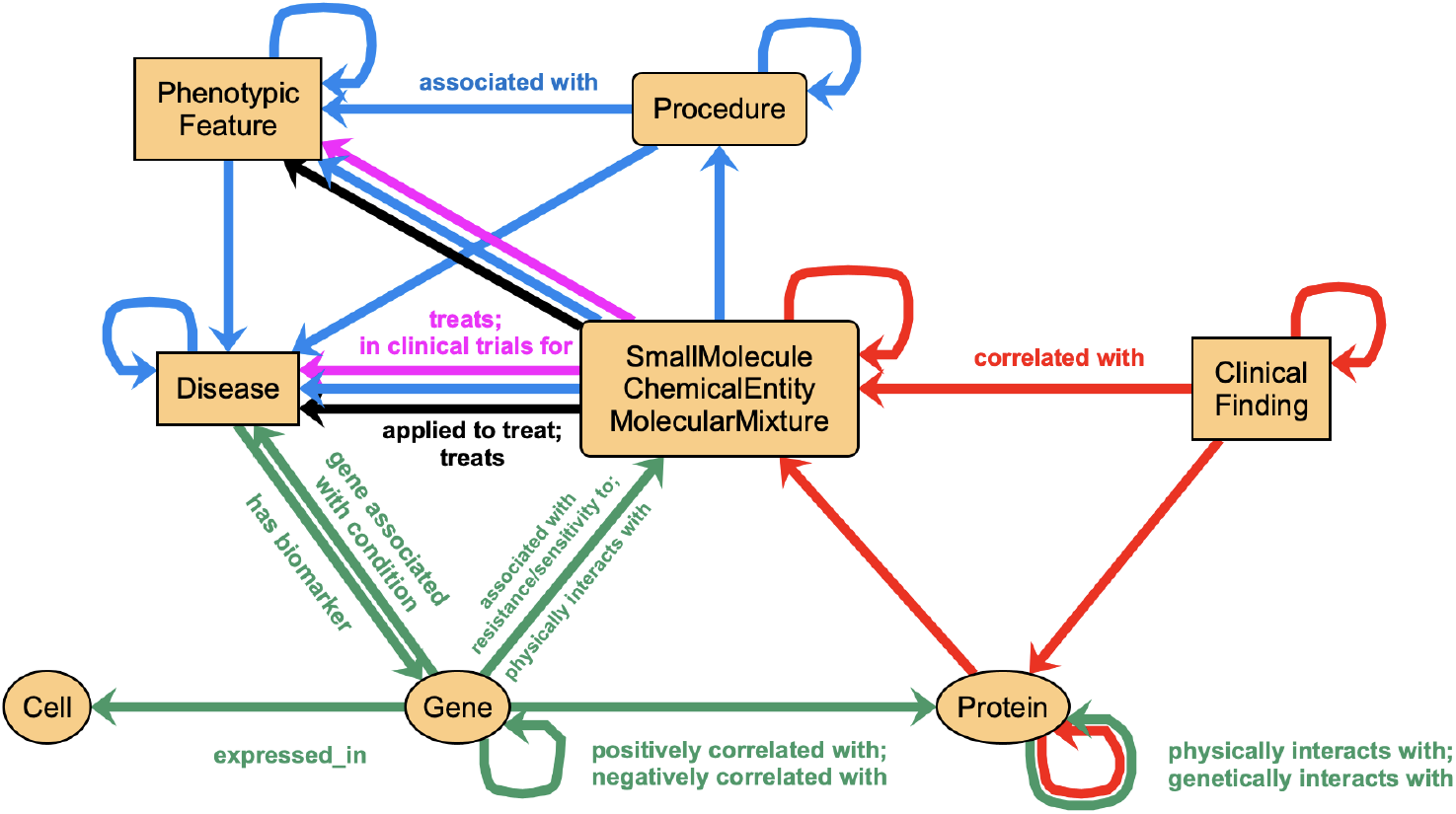
Joint metagraph of the KGs described in this manuscript. Cyan: Clinical Connections, all edges use Biolink predicate “associated with”. Magenta: Clinical Trials, edges use Biolink predicates “in clinical trials for” and “treats”. Black: Drug Approvals, edges use Biolink predicates “treats” and “applied to treat”. Green: BigGIM-DrugResponse, edges as labeled. Red: Wellness Multiomics, all edges use Biolink predicate “correlated with”. See Methods for the definitions of the Biolink classes used. For simplicity, in this figure we depict together the three Biolink classes ‘SmallMolecule’, ‘ChemicalEntity’, and ‘MolecularMixture’.

### 2.2. BigGIM-DrugResponse KG Hub

BigGIM-DrugResponse KG integrates several smaller KGs encompassing diseases, genes/proteins, and drugs/chemicals by statistical modeling and machine learning from large public datasets, as well as collecting publicly available knowledge resources. As such, it may be considered a “KG hub”. BigGIM-DrugResponse KG includes assertions about drug-target and gene-gene interactions, disease-gene associations, and drug-gene associations, focusing on response and resistance to drugs (**Table 1**). The genes are quantified at the levels of genetic variants, gene expression, and protein level. Knowledge assertions are derived from public knowledge resources or learned from specific contexts such as different disease types, tissue types, and patient cohorts through statistical approaches. Multiomic cancer data sources include The Cancer Genome Atlas (TCGA)^24,25^, Genomics of Drug Sensitivity in Cancer (GDSC)^25,27^, and gene expression in normal tissues from the Genotype-Tissue Expression (GTEx) project^23^.

### 2.3. Clinical Trials KG

The basic question that may be asked of this KG is, “what interventions have been clinically evaluated for a given condition/disease?” Clinical Trials KG encodes assertions derived from the descriptions of clinical trials in the ClinicalTrials.gov database. We created a KG derived from the clinical trials dataset Aggregate Analysis of ClinicalTrials.gov (AACT)^28^. AACT includes information about each study registered with ClinicalTrials.gov, including protocol information, disease or condition investigated, study design, subject eligibility criteria and/or exclusion, and outcomes.

A total of 514,498 clinical trials were available for extraction as of November 3, 2024, of which we modeled 115,086 based on their characteristics. We used Babel^29^ to map the interventions and conditions to 22,337 biomedical concepts. These concepts are clinical and/or biological ideas, with a distinct meaning given by reference to a terminology or vocabulary identifiable by unique ID or code (CURIEs). Nodes at this stage consist of five types in Biolink Model: interventions, which are represented by the SmallMolecule (n=957), ChemicalEntity (n=6,525), or MolecularMixture (n=35) classes, and the conditions/diseases for which the interventions were tested, represented by the Disease (n=13,045) or PhenotypicFeature (n=1,775) classes. Edges (n=176,656) are expressed using the “biolink:in_clinical_trials_for” predicate. Additional edges (n=13,450) using the “biolink:treats” predicate are generated when an intervention is being evaluated in the context of a Phase 4 trial (after FDA approval). Edges are further annotated with multiple attributes, including trial phase, status, cohort size, age range for eligibility, etc.

### 2.4. Drug Approvals KG

The basic question that may be asked of this KG is, “what interventions have been approved for a given condition/disease?” Drug Approvals KG encodes assertions derived from the product indications in the DailyMed database^15^. Since the drug indications are provided as descriptive, long-form, unnormalized texts as provided by the product manufacturers, and frequently contain mentions of non-indicated conditions (including side effects and contraindications), we identify the most likely target(s) of the indication by cross-referencing with indications provided in the FDA’s Adverse Event Reporting System (FAERS)^16^ database.

As of November 1, 2024, DailyMed included information on labels for 152,812 products. FAERS included information on 20,407,479 adverse event reports (dated from the first quarter of 2004 and through the third quarter of 2024, inclusive), from which we extracted 35,571,841 non-redundant assertions on prescribed treatments and their associated indications. We used Babel^29^ to map the interventions and conditions to biomedical concepts. We then cross-referenced the content extracted from the two sources (DailyMed and FAERS) to determine the approval status of interventions for their reported indications. Based on this classification, we generated 4,117 edges connecting 919 subjects (approved drugs) to 847 objects (indicated conditions), using “biolink:treats” as predicate. We similarly generated 92,056 edges connecting 1,059 drugs to 5,828 conditions, using “biolink:applied_to_treat” as predicate to indicate absence of formal approval for these indications.

### 2.5. Clinical Connections KG

Electronic health records provide potentially rich, but realistically sparse, longitudinal histories of patient data. Researchers have devised methods for extracting clinical and biological research value from these datasets, in the form of distributions and co-occurrence of clinical features such as medications, adverse drug events, phenotypes, risk assessment, outcomes, etc., amongst and between diseases, whilst maintaining patient confidentiality and privacy^30,31^.

Clinical Connections KG encodes connections among diseases, laboratory results and medication orders observed per patient at Providence Health & Services and affiliates (PHSA), which is an integrated healthcare system which serves patients in 51 hospitals and 1,085 clinics across seven US states: Alaska, California, Montana, Oregon, New Mexico, Texas, and Washington. We use logistic regression machine learning (ML) on EHR medical data to develop multivariable classification models. This approach constructs a single large KG, with directed edges, linking baseline factors (patients’ conditions from the past year) to specific outcomes, including common chronic diseases and rare diseases.

We constructed a KG derived by performing 148 logistic regression models on EHR data from years [01/01/2008 - 05/01/2024]. Nodes consist of medical concepts from the EHR including age, sex, conditions, medications, and labs. These were mapped to CURIEs using the OMOP CDM. Nodes were categorized by the following Biolink Model classes: Disease, PhenotypicFeature, ChemicalEntity, and Procedure. Edges are expressed by one of the opposite Biolink Model predicates: “associated with increased/decreased likelihood of”. Edges between these nodes are annotated by attribute types, including provenance, the analysis method used (logistic regression), the AUCROC, p-value, log odds ratio, 95% confidence interval, sample size with the condition, and sample size without the condition.

### 2.6. Wellness Multiomics KG

The Wellness Multiomics KG encodes extensive phenotyping of the Institute for Systems Biology’s (ISB) Wellness cohort^32^, expanding on ISB’s original wellness study of 108 individuals^26^ and integrating many data types including genetics (whole-genome sequencing and/or SNP genotyping), clinical blood tests, salivary cortisol, weight and body-mass index (BMI), blood pressure, health assessments, gut microbiome, blood metabolomics, blood proteomics, activity tracking, sleep tracking, and heart rate.

We have analyzed the ISB Wellness dataset, which represents extensive phenotyping of ISB’s Wellness cohort, and affords many types of correlations and connections to be uncovered. This deep phenotyping data set integrates many data types including genetics (whole-genome sequencing and/or SNP genotyping), clinical blood tests, salivary cortisol, weight and body-mass index (BMI), blood pressure, health assessments, gut microbiome, blood metabolomics, blood proteomics, activity tracking, sleep tracking, and heart rate. The cohort includes 4,879 individuals with at least one blood draw. Integrative analysis of this multidimensional data set has already led to significant novel findings, e.g., on the connection between blood metabolites and the microbiome^32^ and how this reflects on aging^33^.

Based on this dataset, we have created and deployed the Multiomics Wellness KG. This KG includes pairwise correlations among clinical labs, metabolites and proteins within the ISB Wellness dataset. The resulting KG includes 679,420 statistically significant correlations involving 101 clinical labs, 264 proteins, and 830 metabolites, under 27 different stratification modes (see Methods).

### 2.7. Use case

A researcher has an overarching question: “What might be some new ideas for treating type 1 diabetes (T1D)?” or one of many other imaginable biomedical questions. A formal or informal step in the path to an answer is to create a subgraph from a larger KG (or federation of KGs and other knowledge sources) that contains all nodes and edges relevant to a particular form of logic chosen to answer the question. Indeed, if such a subgraph is sufficiently intuitive, then the subgraph itself may serve as the ‘answer’ to the question. We show in this use case that disparate knowledge from federated KGs, including the five KGs presented here, can be used to create subgraphs useful for responding to biomedical queries. A key to providing a pathway to deeper insights is to have disparate types of data considered as part of the reasoning process^34,35^, which can be evaluated with a metagraph such as shown in **Figure 2**.

To illustrate, we show how such a subgraph might be created to address the diabetes question in the preceding paragraph. The researcher (human, or possibly machine) may break the question down into a general two-step approach for reasoning: (1) first, identify mechanisms that cause or influence diabetes, and then (2) identify interventions that influence those causes. At this level of logic, it becomes natural to leverage a KG for reasoning, both to identify mechanisms and to identify interventions. There are many approaches to identify mechanisms; for the purposes of this vignette we mention three: (a) identify an existing drug known to treat diabetes, and conclude that the molecular physiological subsystem it influences also influences diabetes, (b) identify a candidate drug posited to treat diabetes, and conclude that drug is a candidate because an expert hypothesizes that the molecular physiological subsystem it influences also influences diabetes, (c) identify knowledge in the KG that connects a molecular subsystem to diabetes. There are many approaches to identify interventions; for the purposes of this vignette, we mention three: (i) identify drugs or lifestyle interventions that target identified molecular subsystems, (ii), identify drugs that are bioinformatically similar to other drugs that target identified molecular subsystems, and (iii) identify drugs already known to target diabetes. The various categories of reasoning mentioned are neither necessarily mutually exclusive nor guaranteed to be error proof; rather, they represent an automated approach to hypothesis generation. A full review of KG reasoning is outside the scope of this article; Chen et al.^36^ provide an entry into this literature, including deductive, inductive, and abductive reasoning. In **Figure 3**, we show a simple subgraph designed to capture some of the elements that could be used to support logic through the above approaches. **Figure 3** is designed to show only *some* example nodes and edges; in practice, most power users would create a subgraph with many more nodes and edges to enable stronger logical inferences. **Figure 3** is constructed to provide a subgraph that supports reasoning to produce the already empirically known result that teplizumab can treat or prevent the progression of T1D^37^, and could be used for logic related to understanding that empirical result (possibly reproducing the original inspiration for experimentation^38^) and/or extending that understanding to identify new drugs that might be used based on similar reasoning upon KG knowledge.

**Figure 3.**
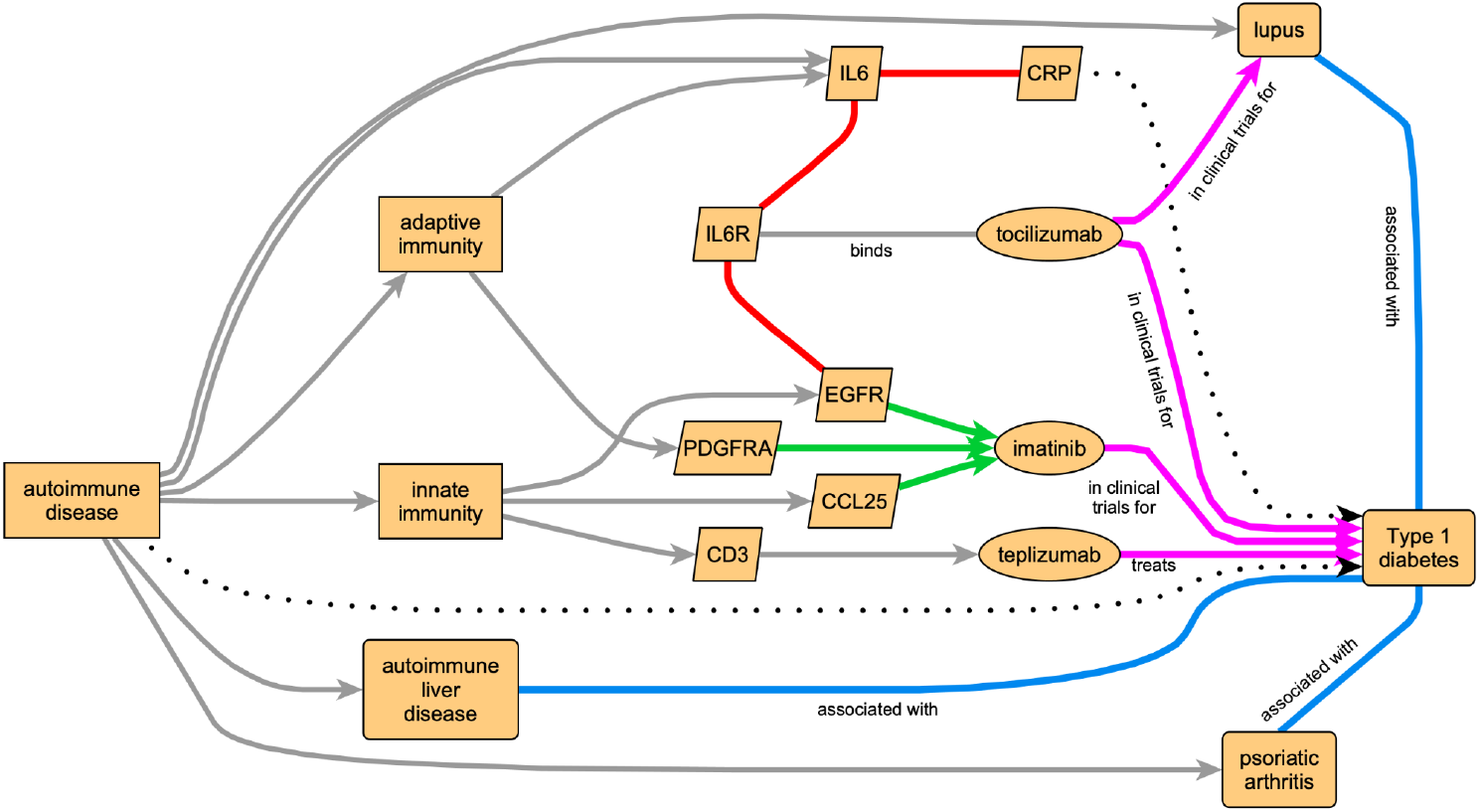
Combining knowledge from multiple diverse KGs can lead to integrative insight in biomedical research. In this example use case, leveraging KGs to investigate causes and treatments for type 1 diabetes (T1D), permits both obvious logical inferences (such as “T1D is an autoimmune disease”) and other inferences such as potential drug treatments for T1D. Colored arrows denote edges sourced from our various KGs (cyan: Clinical Connections; green: BigGIM-DrugResponse; red: Wellness Multiomics; magenta: Clinical Trials), as in **Figure 2**. Gray arrows denote edges from other Translator KGs. Dotted arrows represent inferences.

Some of the intermediate reasoning steps or conclusions that might be supported from reasoning on this or a larger but similarly inspired subgraph might include (1) T1D is an autoimmune disease, (2) treatments influencing both adaptive and innate immunity may be useful in treating autoimmune diseases, (3) C-reactive protein (CRP) is a biomarker for some autoimmune disease and might be a biomarker for others, (4) some drugs targeting a particular autoimmune disease might also be useful for treating another autoimmune disease, (5) many autoimmune diseases are associated with reduced CD3 counts, presumably in response to therapy, and (6) therapies that reduce CD3 counts might work in T1D. Types of logical inferences from KGs may be general, such as “Treatments targeting both innate and adaptive immunity may benefit T1D” or be specific, such as, “Toclizumab might benefit early stage T1D”. Inferences may be based on many underlying edges, each contributing a small amount to a conclusion (e.g., “T1D is an autoimmune disease”) or may be based on one or few edges (e.g., “Teplizumab treats T1D”). Inferences may require “single hop” or multistep “multiple hop” logic on the KG. A general single-hop inference might be: “inflammation causes T1D”. A specific multi-hop inference might be “defects in NLRP3 impair inflammasome function, decrease inflammation, and therefore protect against T1D”. The generality and complexity of inquiries to a KG should depend on the needs of the user and the richness of the data pertinent to the inquiry stored in the KG. A user interface that has customizable parameters and constraints, such as that provided by Translator at https://ui.transltr.io/, can facilitate scoping in or out to find useful subgraphs to motivate and support further investigations.

## 3. Discussion

Many types of biological data can be transformed automatically into knowledge, stored in KGs, and analyzed integratively using systems like the Biomedical Data Translator. KGs provide an important source of information for human or automated biomedical and translational reasoning. There is increasing recognition of a need for automated extraction of knowledge from data. Thanos et al.^39^ describe a framework for “knowledge creation based on the exploitation of the knowledge hidden in huge data volumes of research data”, including (1) creation of linked information spaces; (2) services to make these information repositories discoverable, accessible, understandable and reusable; (3) navigational services; and (4) workflow services providing APIs and user interfaces. Translator provides an infrastructure that integrates all these features. KGs provide a fundamental base of knowledge underlying the Translator ecosystem. In this paper, we have documented how five biomedical KGs were inspired and produced. These KGs can be used stand alone, as part of Translator, or as part of any federated reasoning system (e.g., KGAREVION^40^) capable of drawing upon knowledge stored in KGs.

There is considerable prior art in the generation of KGs from EHRs. A full review is beyond the scope of this paper. Notably, Morris et al. created Scalable Precision Medicine Oriented Knowledge Engine (SPOKE)^41^. Notably SPOKE also automatically generated knowledge with unsupervised machine-learning to create vectors encoding the importance of each EHR code^42^. Liu et al. used ontology mapping and natural-language-processing (NLP) to create a rare-disease KG (OARD)^43^. Rotmensch et al. automated a workflow to link diseases and symptoms from EHR^44^. Chandak et al. developed PrimeKG which integrates multiple resources that describe diseases with relationships representing ten major biological scales^45^. Santos et al. developed a Clinical Knowledge Graph (CKG) focused on proteomics data^46^. Our present work provides additional approaches to knowledge generation and KG creation, expanding the ecosystem of biomedical KGs.

Production of a KG may require addressing specific challenges, including handling large datasets, ensuring privacy, selecting appropriate cohorts, managing data errors and missing values, and achieving concept standardization. BigGIM required considerable compute infrastructure to process very large data sets. Wellness KG required considerations of privacy, cohort selection, and concept mapping. The Clinical Trials and Drug Approvals KGs required concept mapping and data cleaning for user-entered data. The Clinical Connections KG required considerations of differential privacy, computational expense, need for domain knowledge, missingness, and mapping associations based on implicit knowledge. Such challenges should be documented as appropriate in individual KG nodes or edges, but also in overall published methods for KG construction.

Each of the edges in the Wellness Multiomics KG encodes a correlation based on a particular stratification (subpopulation) of the data, as described above. In many cases these stratifications represent ordered classes (such as age ranges: “less than 35”, “between 35 and 55”, “more than 55”), and in some cases unordered classes (“male”, “female”). These may aid in certain types of logical inference, particular mechanistic or causal inference, that may leverage dose-response logic. An alternative data structure for encoding these stratified edges would be to provide each as an annotation to a single KG edge; however, that would defeat the efficiency of many potential downstream graph-based algorithms. An example of such stratification-empowered detail is the relationship between CRP and IL6 (**Figure 3**). Individuals who never use alcohol have a correlation of 0.60; those with daily alcohol use have a correlation of 0.53. The range of correlation is even greater across populations, from 0.57 (Hispanic/Latino) to 0.70 (South Asian). These KG edges and logic performed on them suggest that specific relationships between immune subsystems depend on both environment (alcohol use & culture) and genetics (genetic ancestry). A conclusion might be that immunomodulatory therapies directed at autoimmune diseases such as T1D should take into account such influences in a personalized approach to medicine.

KGs are a natural fit for many types of biological data and reasoning—particularly across biological pathways and networks. KGs are compact, human understandable, and machine interpretable. Lots of algorithms can perform computations on KGs, and there is an active coding community (e.g., Neo4j). Trees are a form of graph, so many graph algorithms and tree algorithms are similar—if not identical. However, there are limitations to KGs in general, as well as to specific KGs. For example, certain types of knowledge are difficult to store in KGs, including conditional logic and multi-step conditional dependencies (e.g., the citric acid cycle). These issues can be addressed by using different types of knowledge storage (e.g., relational databases) within a larger federated ecosystem, such as Translator. Ultimately, we predict that the best reasoners (such as Translator) will reason on knowledge stored in multiple different types of data structures, including KGs. Within each specific KG, there are tradeoffs between choices of ontologies, context (e.g., adult or pediatric), resolution/scale (e.g., encoding SNPs or genes or both as nodes), and many other choices that also should be reflected in node and edge provenance, as well as overall published methods for KG construction. All KGs should be validated before deployment for internal consistency, faithful representation of underlying data, and correct knowledge inferences from data. However, this does not guarantee the utility of a particular KG for a particular purpose; additional use-specific (e.g., drug repurposing) validation should be performed for any critical end-use. Many biomedical knowledge graphs have many edges and multitudes of paths between distant nodes. Many of these paths are distracting due to the low quality (i.e., high uncertainty) of many edges in most KGs. Scoring these paths is beyond the scope of this manuscript, but approaches include excluding low quality edges, employing better quality metrics, and logic algorithms robust to distractions.

Human expertise should never be lost, even as automated intelligence tools are increasingly leveraged. Currently, most automated knowledge generation tools used to populate KGs are simple. Algorithms are chosen because they are straightforward and parsimonious. More complex and nuanced insights based on complex chains of reasoning may be missed by automated tools^22^. Large sets of data can be transformed into lossy KGs by these simple tools. These can be very useful, but may fail certain uses. Therefore, even as large sets of raw data are transformed into KGs, these raw data should not then be discarded or disregarded—they might be useful for certain nuanced investigations, including some types of hypothesis-driven inquiry. If possible (e.g., allowed by privacy considerations), multiple KGs can be constructed from the same data set using different choices and parameters for extracting knowledge from data. Increasingly sophisticated knowledge extraction is a future of KG construction. Large-language model (LLM) based artificial intelligence (AI) algorithms were not used in the creation of knowledge for the KGs described here, but are and will be used for generating a next generation of biomedical KGs.

Theories of knowledge and the relationship of data to knowledge date back to the earliest philosophical discourses, such as Plato’s *Theaetetus*. Modern informatics has driven an urgency for more precision and uniformity in definitions of specific concepts such as “data” and “knowledge”, but uniformity of definitions has yet to be implemented. However, even in the current epoch of unclear or fuzzy boundaries between the concepts of ‘data’ and ‘knowledge’, a hierarchy of knowledge can be conceptualized, ranging from atomic bits of data to deeply profound integrative understanding held in common by most human minds. Each KG represents a particular level (or may bridge several levels) in this hierarchy of knowledge. Of the KGs presented here, some reflect basic elements of data—with minimal transformation. For example, Drug Approvals and Clinical Trials KGs largely reflect a Boolean value of whether or not a drug has been approved or simple ‘drug is in trial’ triple. Knowledge added is largely limited to checks for validation, consistency, and coherency. BigGIM-DrugResponse, Clinical Connections, and Wellness Multiomics KGs climb this hierarchy higher, using machine learning to produce ‘knowledge assertions’ or outright knowledge not readily apparent in large raw datasets. Other KGs may include even higher levels of knowledge, such as facts generated by advanced artificial intelligence or extracted by human readers from highly acclaimed and time-tested review articles. Whether any given KG truly enshrines knowledge depends to some extent on continuing evolving definitions of ‘knowledge’. However, all KGs—including the five presented here—are steps on the hierarchy of enlightenment, and can aid basic and translational biomedical research.

## 4. Methods

Our data-to-knowledge pipeline starts with data extraction, transformation, and loading (ETL) into intermediary data structures, followed by statistical analysis and mapping to relevant ontologies and to the Biolink Model. Biolink provides an open-source data model that formalizes relationships between biomedical data structures^10^. The resulting KGs consist of nodes (concepts) linked by edges (relationships). The nodes represent well-identified biomedical concepts, mapped to terms from suitable ontologies. These ontologies encompass diseases, genes, clinical measurements, and other biomedical concepts. The edges represent relationships between concepts encoded in the nodes. Both nodes and edges are annotated with ‘attributes’ that provide additional information beyond their label, including links to databases and descriptive information as well as provenance (**Figure 1**).

### 4.1. General overview of KG development

The framework for our pipeline to generate and deploy KGs consists of (1) ETL/extraction, (2) statistical analysis, (3) semantic standardization through Biolink modeling^10^, (4) KG generation, and (5) deployment as an open source web service that implements a standard API. In addition to the semantic standardization, each KG can be represented as a pair of Knowledge Graph Exchange (KGX) tab-separated value (TSV) text files^47^ with nodes and edges, which permits export to other programs such as Cytoscape^48^ (cytoscape.org) and facilitates quality validation, as described below.

#### Data standardization across KGs and the Translator ecosystem

For reasoning to occur on a concept, a human or machine reasoner must make connections between multiple data incorporating that concept. This requires the reasoner to recognize that distinct data refer to the same concept. In a data universe awash in synonyms and parasynonyms, a good KG must standardize synonymous concepts with ontology-derived nomenclature. Depending on the coarseness of these ontologies and data sources, parasynonyms must also be mapped to ontology terms with nearby meanings. Multiple KGs providing information to a reasoner should unify these nomenclature mappings; if they did not, the burden of recognizing synonyms would fall upon the reasoner—which would be inefficient. For the KGs discussed here, nodes were mapped to names from standard ontologies including: NCBIs Gene (www.ncbi.nlm.nih.gov/gene) & HUGO Gene Nomenclature Committee (HGNC) (www.genenames.org) for genes; UniProt (www.uniprot.org) for proteins; Pubchem (pubchem.ncbi.nlm.nih.gov), Chemical Entities of Biological Interest (ChEBI) (www.ebi.ac.uk/chebi), the Human Metabolome Database (HMDB)^49^, Chemical Abstracts Service (CAS) (www.cas.org), Kyoto Encyclopedia of Genes and Genomes (KEGG)^50^, RefMet^51^, & Experimental Factor Ontology (EFO)^52^ for chemicals and metabolites; Drugbank (go.drugbank.com) & Chemical Database of Bioactive Molecules with Drug-Like Properties (ChEMBL) (www.ebi.ac.uk/chembl) for drugs; Mondo Disease Ontology (MONDO) (mondo.monarchinitiative.org) for diseases; Logical Observation Identifiers Names and Codes (LOINC) (loinc.org) for clinical labs, & Human Phenotype Ontology (HPO) for clinical labs outside of reference range. Nodes with synonymous names from multiple ontologies were assigned a single unified (preferred) compact universal resource identifier (CURIE), as specified in Biolink Model^10^. Each node is identified with a single unified CURIE using Babel^29^, a relational database underlying Translator’s Name Resolver/Node Normalizer (name-resolution-sri.renci.org/docs). Additional synonyms were annotated to that node along with other annotations including those necessary for provenance. For clinical KGs, for the analytes that have significant correlations with other analytes, we performed a detailed manual curation of LOINC codes for the attributes in the chemistries table, and modified the Biolink concepts of a subset of nodes to ClinicalFinding to retain LOINC codes that best preserve the identity of a node. Any nodes that failed to map to any CURIE were excluded from the KGs, as they could not be called by the Translator API. At this time, the Translator ecosystem allows gene and protein concepts to be conflated; gene names are synonymous with their gene product (e.g., proteins); the unified concept is primarily identified by its NCBI Gene symbol.

#### Definitions of Biolink Classes represented in our KGs

ChemicalEntity: physical entity that pertains to chemistry or biochemistry. SmallMolecule: molecular entity characterized by availability in small-molecule databases of SMILES, InChI, IUPAC, or other unambiguous representation of its precise chemical structure; for convenience of representation, any valid chemical representation is included, even if it is not strictly molecular (e.g., sodium ion). ClinicalFinding: this category is currently considered broad enough to tag clinical lab measurements and other biological attributes taken as ‘clinical traits’ with some statistical score, for example, a p value in genetic associations. Disease: disorder of structure or function, especially one that produces specific signs, phenotypes or symptoms or that affects a specific location and is not simply a direct result of physical injury. A disposition to undergo pathological processes that exists in an organism because of one or more disorders in that organism. Drug: substance intended for use in the diagnosis, cure, mitigation, treatment, or prevention of disease. Gene: region (or regions) that includes all of the sequence elements necessary to encode a functional transcript. A gene locus may include regulatory regions, transcribed regions and/or other functional sequence regions. MolecularMixture: chemical mixture composed of two or more molecular entities with known concentration and stoichiometry. PhenotypicFeature: combination of entity and quality that makes up a phenotyping statement. An observable characteristic of an individual resulting from the interaction of its genotype with its molecular and physical environment. Protein: gene product that is composed of a chain of amino acid sequences and is produced by ribosome-mediated translation of mRNA.

### 4.2. BigGIM-DrugResponse KG

BigGIM-DrugResponse KG connects diseases, genes, proteins, and drugs or chemicals by statistical and machine learning modeling on large public datasets, as well as including knowledge from publicly available resources. Data and knowledge sources include: protein-protein interactions from the human reference protein interactome^53–58^ and BioGRID^59^; drug-target interactions in DrugCentral^60^, Therapeutic Target Database^61^; text mining of scientific literature; genetic interactions from Biogrid^62^; gene-gene coexpression; disease associated genes; gene - drug response relationships; and cell type - gene signatures relationships.

#### Disease associated gene/proteins

Disease-gene associations highlight genes that are highly frequently mutated in each tumor type (> 5% samples, and mutated in at least 5 samples in the TCGA dataset), and at the same time, those genes that have been predicted as cancer driver genes in the literature^63^. We also included reported disease associated variants for other diseases^27^. The disease-gene edges also provide cancer type specific gene expressions extracted from cancer cell lines. We extracted cancer type specific highly expressed genes from the gene expression data from GDSC^25,27^. We compared the cell lines for each cancer type to the other cancer types using T-test followed by Benjamini–Hochberg multiple testing correction. Effect size was measured to quantify the difference of gene expression between one cancer type vs others. Genes that show significant up-regulation in one cancer type compared to the other cancer types were selected (FDR < 0.05, Effect size > 0). We further ranked the expression of one gene for all the GDSC samples, and got the rank of one gene expression from samples in one cancer type. We filtered genes with median rank greater than 0.75 as the final marker genes for each cancer type.

#### Gene-gene interaction extraction

We measured co-expression with Spearman correlation coefficients and defined, as positively correlated or negatively correlated gene pairs, those with correlation coefficient > 0.5 or < -0.5, with p-value (Benjamini-Hochberg adjusted) < 0.05. For tissue-specific gene co-expression analysis, we used Genotype-Tissue Expression (GTEx) project data (version 8) to determine the co-expression of two genes in different tissue types. For tumor-type specific gene co-expression analysis, we used TCGA Pancancer Atlas data. We collected protein-protein interactions from BioGRID and from The Human Reference Protein Interactome Mapping Project (HuRI/HI-union) (www.interactome-atlas.org)^53^. BioGRID includes physically interacting gene products and genetically interacting genes^59^.

#### Drug Response extraction

We focused on Gene (expression) associated with sensitivity to Drug (Small Molecules). Drugs with IC50 <= 0.5 µM in at least 3 cell lines in one cancer type were used for analysis. We defined the resistance group and sensitive group of cells using the threshold of 1^st^ quartile and 3^rd^ quartile: sensitive group: IC50 < 1^st^ quantile (IC50), and resistant group: IC50 > 3^rd^ quantile (IC50). Student T-test was used to compare the gene expression values between the resistant group and the sensitive group, followed by Benjamini-Hochberg adjustment. Results with FDR < 0.25 were selected and presented in the API. We also analyzed the gene mutation that may alter the drug sensitivity by testing the difference of drug sensitivity by comparing the wild-type group and the mutated group. Drugs with IC50 <= 0.5 µM in at least 3 cell lines in one cancer type were used for analysis.

#### CellMarker extraction

CellMarker interactions were constructed from the CellMarker 2.0 human cell types table^64^. The table was converted to a csv file by treating the Gene as the subject and the Cell Type as the object, with an expressed_in relation. Only entries with both NCBI Gene IDs and Cell Ontology IDs were included. UBERON identifiers for the tissue types were used.

### 4.3. Clinical Trials KG

Clinical trials evaluate the effectiveness of interventions—including lifestyle changes, procedures, and medications—on clinical conditions (diseases). The ClinicalTrials.gov registry solicits collection of protocol information and result summaries for registered studies^65^. We extracted data on interventions and their target conditions from the Aggregate Analysis of ClinicalTrials.gov (AACT) database^28^. The Clinical Trials KG currently transforms the information in the registry into a convenient KG format, but does not generate new knowledge from data.

#### Data selection and processing

We obtained from AACT information on interventions, conditions, and related tables from 514,498 studies (access date: November 3, 2024) and merged the tables using the National Clinical Trial identifier (NCT ID) as shared key. We select those studies for which the “primary purpose” field is either “Treatment” or “Prevention”; which have at least one intervention of type “Drug”, “Biological”, “Dietary Supplement”, or “Combination Product”; and which have at least one stated condition.

#### KG building

Core triples consist of a node representing an intervention (chemical compound, drug, procedure, or other therapies) to a node representing a condition, via an edge annotated with the NCTID of the clinical trial where that intervention was tested for that condition. Nodes are categorized into Biolink classes using Babel^29^. We classify each listed intervention by whether it is used in experimental arms, in control arms, or in both. For interventions used solely in experimental arms, an edge is created with it as ‘subject’, “biolink:in_clinical_trials_for” as ‘predicate’, and each condition as ‘object’. If more than one intervention is used in experimental arms, the resulting edges are annotated to indicate the reduced confidence level in the assertion that the interventions were tested for the conditions. If there are multiple edges with the same subject, predicate, and object, they are combined into a single edge that collects details about all studies that yielded such edges. Finally, if a high-confidence edge exists with maximal study phase of 4, an extra edge is generated with the same subject and object, with predicate “biolink:treats”, and with the supporting data pertinent to the underlying phase 4 trial(s).

### 4.4. Drug Approvals KG

Information on FDA drug approvals can be obtained from DailyMed^15^, which includes structured product labels (SPL) in parsable extensible markup language (XML) format. The drug indications are provided as descriptive, long-form, unnormalized texts as provided by the product manufacturers, and frequently contain mentions of non-indicated conditions (including side effects and contraindications). We implemented a procedure for identifying the most likely target(s) of the indication by cross-referencing with indications provided in the FDA’s Adverse Event Reporting System (FAERS)^16^ database.

#### Data selection and processing

We obtained from DailyMed information on labels for 152,812 products (access date: November 1, 2024) as zip files spanning the prescription (RX), over-the-counter (OTC), homeopathic, animal use, and remainder sections. We extracted from the zip files SPL descriptions in XML format, and then identified in each product’s XML file information about new drug approval (NDA) codes, active ingredients, indications, and boxed warnings. We separately obtained from FAERS information on 21,964,449 adverse event reports (dated from the first quarter of 2004 and through the third quarter of 2024, inclusive); dropped cases as indicated; de-duplicated the cases based on identifiers, retaining for each case the most recent report, which yielded a final list of 13,995,777 unique cases; extracted the information on prescribed treatments and their associated indications, yielding 35,571,841 assertions connecting drugs to indications, which we mapped to CURIEs using Babel^29^. Finally, we used the names of the indications in FAERS, and any of their synonyms, to evaluate their presence in the DailyMed-extracted indication texts.

#### KG building

Core triples consist of a node representing an intervention (drug, supplement) to a node representing a condition, via an edge annotated with relevant NDA and SPL codes from DailyMed, and the number of supporting unique FAERS cases. Nodes are categorized into Biolink classes using Babel^29^. For interventions matching between DailyMed and FAERS, an edge is created with the intervention as subject, “biolink:treats” as predicate, and the indicated condition as object. For interventions linked to indicated conditions in FAERS but without a matching approval in DailyMed, an edge is generated with the treatment as subject, “biolink:applied_to_treat” as predicate, and the indicated condition as object.

### 4.5. Wellness Multiomics KG

The Wellness Multiomics KG generates knowledge from data by extracting relationships from dense data collected on a large cohort of well individuals.

#### Data selection and processing

We included in this study the ‘chemistries’, ‘metabolomics’, and ‘proteomics’ tables in the ISB Wellness dataset^26^, version snapshot May 31st, 2019. These tables respectively include data on clinical labs, metabolites, and proteins. To focus on healthy individuals, we excluded from analysis any individuals for which disease-specific assays were performed, as these individuals were likely to not be well (i.e., they had a disease). We further excluded all clinical chemistries labeled as ‘reflexive’ in the dataset; ‘reflexive’ testing is blood testing that was performed depending on whether the previous test result for the same analyte was out of range, and inclusion of reflexive tests leads to biases. We retained only baseline data for each individual, before any wellness interventions. *Cohort stratification:* We stratified the cohort based on multiple self-reported demographic and lifestyle parameters. These included age range (using 35 years old and 55 years old as cutoffs), self-reported biological sex, self-reported race/ethnicity, alcohol use frequency, tobacco exposure status, marijuana use, other illicit drug use, and family structure (number of children). See Supplementary Table 2 for full details. We produced joint analyte tables for the entire cohort of healthy individuals (N=4,234), and separately for each stratification.

#### Data analysis

We computed all-against-all analyte Spearman correlations for the entire cohort and, separately, for each stratification. We used Spearman (rank) correlations since relationships between the analytes are not always linear. For each joint analyte table, we (1) identified analyte pairs where both analytes had non-null values for at least ten individuals, (2) computed a Spearman correlation for each such pair of analytes, (3) considered the count of such pairs as the number of tests performed, (4) retained the resulting Spearman rho and the uncorrected p-value, for analyte pairs where p-value times number of tests was < 0.05. This computation step yielded a list of significantly correlated analyte pairs, and the number of tests performed, for each stratification. Based on the number of resulting correlations, we dropped underpowered stratifications including Middle Eastern ancestry (42 correlations), Native American ancestry (86 correlations), and drug use other than marijuana (57 correlations). We then computed the total number of tests performed across all remaining stratifications (N=1,189,745) and retained Bonferroni-corrected p-values < 0.05 as significant (N=653,226).

#### KG building

We generated a KG for the Wellness dataset, representing each analyte (clinical lab, protein, or metabolite) as a node and each significant correlation as an edge. For the nodes, we used Biolink classes: biolink:ClinicalFinding, biolink:Protein, and biolink:SmallMolecule for clinical labs, proteins, and metabolites, respectively. For the edges, we used biolink:correlated_with, corresponding to RO:0002610 in the Relations Ontology^66^. Each edge is annotated with attributes denoting the statistical test used (Spearman correlation, NCIT:C53236), the effect size estimate (STATO:0000085), the sample size used to compute the correlation (GECKO:0000106), the Bonferroni-corrected p-value (NCIT:C61594), and the stratification used, if any (**Supplementary Table 2**).

### 4.6. Clinical Connections KG

The Clinical Connections KG generates knowledge by extracting relationships from EHR data.

#### Data selection and processing

We analyzed Providence Health & Services (PHS) electronic health records within a secure data enclave, and only exported the final analysis results (nodes and edges, see **Figure 4** for the pipeline). Theoretically, tens of thousands of medical conditions could be included. To keep costs in scope for this proof-of-concept research work, a curated subset of disease concepts were selected by people with medical training, considering chronic conditions with higher relative prevalence in the United States, as well as several rare diseases. Based on medical relevance to the selected conditions, 148 conditions, 366 relevant medications and 115 laboratory tests were included. For conditions/diseases and medications status, we use the EHR-reported status to determine whether a patient has a history of those features. For continuous laboratory results, we use the EHR-reported reference range to determine whether a particular result is high or low. and used LOINC2HPO^67^ to map results to a phenotype (for example, sodium below reference range was mapped to hyponatremia and above reference range was mapped to to hypernatremia). To support privacy, age was binned (0-17, 18-49, 50-74, >= 75 years).

**Figure 4.**
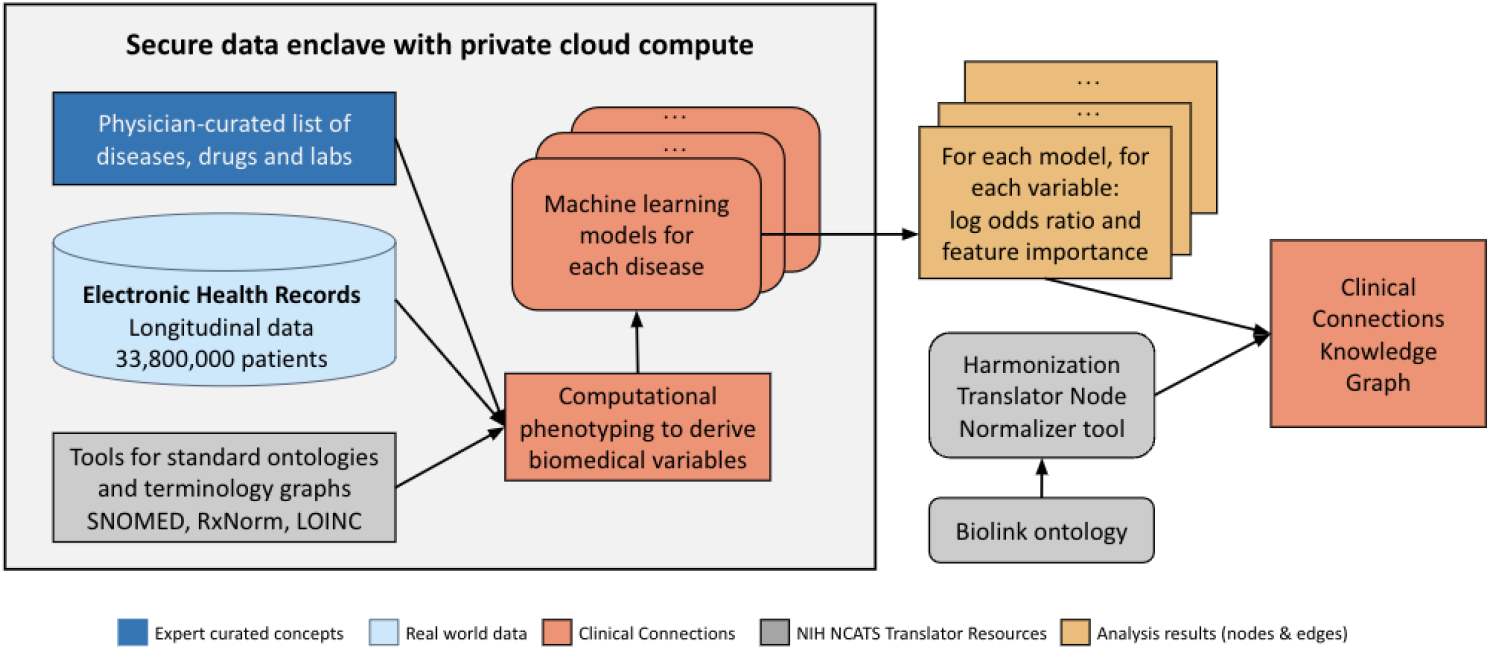
Overview of the workflow to generate the Clinical Connections KG from real world data in electronic health records.

#### Concept mapping

Data were mapped to concepts in the Observational Medical Outcomes Partnership Common Data Model v6.0 (OMOP CDM) in July 2023 by mapping to the Observational Health Data Sciences and Informatics (OHDSI) vocabulary list (athena.ohdsi.org/ vocabulary/ list). SNOMED data was derived from the United States Edition of the SNOMED CT Browser (browser.ihtsdotools.org), and RxNorm data was derived from the NIH RxNav browser (mor.nlm.nih.gov/RxNav). Concepts with less than 10 patients/encounters were excluded to focus on concepts with broader applicability and to protect privacy. Additionally to protect privacy, in the result edge file, patient/encounter count for each concept has been rounded to the nearest hundreds. The nodes of the core-triple for the KG thus constructed consist of the predicted condition/disease from the model, and the independent or feature variable for that model. One of two predicates formalized directed edges between nodes according to the Biolink Model^10^: “associated with increased likelihood of” or “associated with decreased likelihood of”. These edges are annotated with the following information from each model, constituting the attributes of these edges: the AUROC (area under the receiver operating characteristic), p-value, 95% confidence interval, feature importance, feature coefficient, and sample size of patients with and without the condition/disease. The end result: over 39,553 edges representing predictive factors of disease.

#### Model training and concept association analyses

We trained 148 multivariate logistic regression models, with the outcome predicted being one of the 148 conditions/diseases, and independent variables being the combination of those 148 conditions/diseases, 366 medications, 115 lab measurements, and 5 demographic features. Age, sex, and ethnicity were included in all logistic regression models; however, we excluded these demographic features from the resulting KGX files. The Clinical Connections KG uses log odds ratios derived from the logistic regression models to quantify associations between concepts. The AUROC for each model is provided, along with the 95% confidence intervals and p-values. We did not conduct false discovery rate (FDR) adjustment because in the FDR method, p-values are multiplied by m/k (k is the position and m is the number of independent tests) and ranked in an ascending sorted vector of p-values. It potentially rejects true positives among false positives.

### 4.7. Quality Control (QC) and Testing

Steps for evaluating the technical quality and robustness of these KGs are similar. We implemented four layers of quality control, as follows.

#### Preliminary KG evaluation

To evaluate for internal consistency, we implemented a domain-agnostic QC method similar to the method we previously described for BDQC^68^. We perform basic tests of data types and consistency, including checks for repetitions of values & relationships and for missingness on both the node and the edge TSV files. We also evaluate the consistency between nodes and edges: whether declared nodes have no associated edges, and whether edges refer to undeclared nodes. All tests are summarized in a compact JSON format report. We then evaluate any QC flags in the report and make a domain-informed assessment of whether they are justified (e.g., nodes can have no associated edges), or alternatively whether they reflect computational or representation failures (e.g., multiple nodes with identical identifiers, edges linking undeclared nodes). We also used well-known interactions for domain-informed verification. For example, for the BigGIM-Drug Response KG, we use the drug-target interaction as a cross checking of the KG by examining whether the target gene itself is a predictor or biomarker for its targeted drugs. The comparison of evidence such as p-values between the newly generated KG and gold-standard interactions provides an overall metric of confidence.

#### Basic validation

Edges were sampled randomly from KGs and reviewed by subject matter experts for biologic plausibility. For most KGs, ∼40-100 edges are sampled by experts, who also evaluate edge & node metadata including evidence and provenance^69,20^.

#### Internal querying

To perform testing of querying new KGs in-house, we implemented an internal queryable endpoint using fastAPI. Since KGs can be very large, we selected representative edges to assess the integrity of the entire KG. For example, this procedure selected 939 edges out of the 229,614 edges in version 1.3 of the Multiomics Wellness KG. Queries are tested on the internal endpoint to ensure that expected results are returned speedily and to ensure the integrity of the KG. The internal endpoint is similar to a Translator TRAPI endpoint, so passing internal tests minimizes the risks of failures upon external deployment.

#### External querying

KGs are deployed (see next section) for testing by users via virtual knowledge graph interfaces or through the Translator ecosystem (https://ui.transltr.io/). Feedback is then collected from users via Github tickets.

### 4.8. Deployment

KGs available to Translator automated reasoners and to public querying through SmartAPI (smart-api.info)^70^, BioThings (REST API), and TRAPI endpoints. URLs and API documentation can be found at github.com/NCATSTranslator/Translator-All/wiki/ Multiomics-Provider.

#### APIs

We set up BioThings APIs that can be queried directly as REST APIs. API development and deployment consists of the following steps: (a) Generate knowledge from correlation analysis, machine learning model predictions, etc. and structure it in the form of nodes and edge relationships following the KGX format, (b) store the TSV files on a file server, (c) write a parser script and manifest file as described in the BioThings Studio documentation (docs.biothings.io/ en/latest/ tutorial/ studio.html) and store these files in a GitHub repository, and (d) use BioThings Studio to deploy to the Translator’s Service Provider server. A CI/CD flag can be enabled by the Service Provider group for step (d) so that updated KGX files stored on a file server can easily be loaded and merged with the existing KG. Parsers, SmartAPI yaml files, and other utilities are also available in our Github repositories.

#### TRAPI endpoints

We set up TRAPI endpoints that can be queried by Translator’s automated reasoning tools via BioThings Explorer^71^, Plover (https://github.com/RTXteam/PloverDB), and Plater (https://github.com/TranslatorSRI/Plater). The BioThings Explorer tool uses the semantic annotation in a KG’s SmartAPI Registry registrations to check if TRAPI queries may be answerable with knowledge from the KG, to set up queries to a KG’s BioThings APIs, and to transform the responses into the TRAPI standard. The BioThings Explorer tool’s */v1/smartapi/{smartapi_id}/query* endpoint can be used to access KGs individually. Plover is an in-memory Python-based platform designed to host and serve Biolink-compliant KGs as TRAPI APIs; it automatically performs Biolink predicate/class hierarchical reasoning and concept subclass transitive chaining, among other tasks. Plater is a web server powered by Neo4j that exposes Biolink compliant KGs as TRAPI APIs; it automatically performs Biolink predicate/class hierarchical reasoning and concept subclass transitive chaining.

#### Privacy

The Translator Consortium has created novel approaches for hosting clinical data and observational patient data including HIPAA Safe Harbor Plus (HuSH+) clinical data, clinical profiles, Columbia Open Health Data (COHD), and the Integrated Clinical and Environmental Exposures Service (ICEES)^6,30,72,73^. Translator is developed to comply with HIPAA guidelines.

## Supporting information

Supplemental Table 1 - metagraph information

Supplemental Table 2 - wellness graph stratification

## Data availability

Live and updated versions of the Knowledge graphs are available via SmartAPI.

- Clinical Connections KG: https://smart-api.info/ui/eb4e66886fe5c178ae41977cea2c6307
- BigGIM-Drug Response KG: https://smart-api.info/ui/adf20dd6ff23dfe18e8e012bde686e31
- Clinical Trials KG: https://smart-api.info/ui/e51073371d7049b9643e1edbdd61bcbd
- Drug approvals KG: https://smart-api.info/ui/edc04feaf16c12424737988ce2e90d60
- Wellness Multiomics KG: https://smart-api.info/ui/02af7d098ab304e80d6f4806c3527027

## Code availability

The code and instructions to perform the EHR prevalence analysis and other analyses are publicly available on GitHub with no restrictions to access.

- Clinical Connections KG: https://github.com/Hadlock-Lab/clinical_risk_kp
- BigGIM-Drug Response KG: https://github.com/multiomicsKP/drug_response_kp
- Clinical Trials KG: https://github.com/multiomicsKP/clinical_trials_kp
- Drug approvals KG: https://github.com/multiomicsKP/drug_approvals_kp
- Wellness Multiomics KG: https://github.com/Hadlock-Lab/multiomics_wellness_kp

## Author information

### Contributions

GG, JH, GQ conceived of the study. GG, GQ, KN, AJ, SM, MHB, RR, BB, QW, SLG, CX, YY,

AKG, EDM designed and implemented the system. GG, GQ, KN, AJ, AR, YZ, SLG performed analyses. GG, GQ, KN, JCR, QW, JH wrote the manuscript. All authors edited and approved its final version.

## Acknowledgements

We wish to thank Denise Mauldin, Chengzen Dai, Theo Knijnenburg, and John Earls, who contributed coding for early versions of KGs, and Allison Kudla for help with graphic design.

